# A screen of 1,049 schizophrenia and 30 Alzheimer’s-associated variants for regulatory potential

**DOI:** 10.1101/447557

**Authors:** Leslie Myint, Ruihua Wang, Leandros Boukas, Kasper D. Hansen, Loyal A. Goff, Dimitrios Avramopoulos

**Affiliations:** Department of Biostatistics, Johns Hopkins Bloomberg School of Public Health; McKusick-Nathans Institute of Genetic Medicine, Johns Hopkins School of Medicine; Department of Neuroscience, Johns Hopkins School of Medicine; Department of Psychiatry, Johns Hopkins School of Medicine

**Keywords:** schizophrenia, Alzheimer’s, association, gene regulation, reporter assays

## Abstract

Recent genome-wide association studies (GWAS) identified numerous schizophrenia (SZ) and Alzheimer’s disease (AD) associated loci, most outside protein-coding regions and hypothesized to affect gene transcription. We used a massively parallel reporter assay (MPRA) to screen, 1,049 SZ and 30 AD variants in 64 and 9 loci respectively for allele differences in driving reporter gene expression. A library of synthetic oligonucleotides assaying each allele 5 times was transfected into K562 chronic myelogenous leukemia lymphoblasts and SK-SY5Y human neuroblastoma cells. 148 variants showed allelic differences in K562 and 53 in SK-SY5Y cells, on average 2.6 variants per locus. Nine showed significant differences in both lines, a modest overlap reflecting different regulatory landscapes of these lines that also differ significantly in chromatin marks. Eight of nine were in the same direction. We observe no preference for risk alleles to increase or decrease expression. We find a positive correlation between the number of SNPs in Linkage Disequilibrium (LD) and the proportion of functional SNPs supporting combinatorial effects that may lead to haplotype selection. Our results prioritize future functional follow up of disease associated SNPs to determine the driver GWAS variant(s), at each locus and enhance our understanding of gene regulation dynamics.

## Introduction

Over the last decade collaborative genome wide association studies (GWAS) have identified thousands of DNA variants that show robust associations with an array of complex phenotypes (Visscher et al., 2017). For example, over 100 independent loci have been identified contributing to the risk for schizophrenia (SZ) (Pardiñas et al., 2018; Schizophrenia Working Group of the Psychiatric Genomics Consortium, 2014) and based on recent reports from the SZ working group of the psychiatric genomics consortium (PGC, World Congress of Psychiatric Genetics 2017, Orlando, FL) over twice that number of loci will soon be published. In Alzheimer’s disease (AD) a large meta-analysis in 2013 (Lambert et al., 2013) reported 19 risk loci, of which 11 were novel. After completion of this work a larger AD GWAS meta-analysis added 5 more loci. Identifying reliable associations of genetic variants with disease has been a significant first step towards understanding the causes of neuropsychiatric disorders such as SZ and AD. Understanding their functional consequences and linking them to specific affected genes and biological processes is the next necessary step to unlock their translational potential. Across disorders GWAS have taught us some important lessons that help with this goal. For one, variants are most often located in non-coding sequences, and 40% of the time their haplotype blocks do not include coding exons (Hindorff et al., 2009; Manolio et al., 2009; Visel, Rubin, & Pennacchio, 2009). Further, they concentrate in regions of regulatory DNA marked by deoxyribonuclease I (DNase I) hypersensitive sites (DHSs) (Maurano et al., 2012), and Quantitative Trait Loci (eQTL) studies suggest that they are often regulatory (Cookson, Liang, Abecasis, Moffatt, & Lathrop, 2009; Hindorff et al., 2009; Schaub, Boyle, Kundaje, Batzoglou, & Snyder, 2012). It must be noted that such studies likely underestimate how often disease variants are regulatory since variation in the time, place, or circumstances under which a regulatory sequence is activated can lead to false negatives both for regulatory DNA marks and eQTLs.

Typically each GWAS locus contains multiple variants in high linkage disequilibrium (LD), all of which show strong evidence of association. This restricts efforts to identify the one or few variants that drive the association signal at a given locus and to evaluate their functional consequences. For each locus that is associated with disease risk, there must be at least one strongly correlated functional variant explaining the observed association, while it is possible that in some loci there are multiple functional variants in LD with each other along with other “passenger” variants. Such loci might contain specific haplotypes under selective pressure because of their distinct functional profiles, as recently reported for Mendelian disease mutations (Castel et al., 2018). Episomal reporter assays have been commonly used to assess the potential of DNA sequences to drive transcription and to identify functional variants among the many in LD. In these assays, a plasmid is constructed carrying the candidate enhancer sequence inserted next to a minimal promoter driving the expression of a reporter gene. The plasmid is transfected into cells without genome integration and the expression of the reporter gene is measured and compared across different sequences. More recently, to meet the needs of genome level analysis, these types of assays have been redesigned to achieve higher throughput. Examples of such high throughput assays include massively parallel reporter assays (MPRA) (Inoue & Ahituv, 2015) that examine thousands of candidate sequences in parallel and self-transcribing active regulatory region sequencing (STARR-seq) which screen entire genomes for enhancers based on their activity (Muerdter, Boryń, & Arnold, 2015).

Because of our interest in SZ and the large number of reported robust associations we decided to screen by MPRA the many variants in LD at SZ GWAS loci identified by the largest GWAS available at the time of design (Schizophrenia Working Group of the Psychiatric Genomics Consortium, 2014) (36,989 cases and 113,075 controls). We designed our screen to maximize the number of tested variants at the expense of confidence on individual positive results, which will be strong candidates for more elaborate functional testing. In our list of variants of interest and because of some available space on the synthesis array after excluding the largest of the SZ LD blocks, we included a few of the variants reported to be associated with AD by the largest GWAS available at the time of design (Lambert et al., 2013) (17,008 cases and 37,154 controls), another interest of our laboratory.

## Materials and methods

### Selection of variants

Variants were selected from two large GWAS, one for schizophrenia (Schizophrenia Working Group of the Psychiatric Genomics Consortium, 2014) and one for Alzheimer’s disease (Lambert et al., 2013). The schizophrenia PGC2 data was downloaded from https://www.med.unc.edu/pgc/ which provides data on the association between schizophrenia and 9,444,231 imputed SNPs across the genome. We first determined the size of the groups of SNPs that were in LD with each other and therefore represented the same association signal. To maximize the number of independent loci investigated in our assay while comprehensively examining each tested locus, we followed specific selection rules. First, we identified the lead SNPs at each locus and identified all SNPs in the same locus with p-values up to 15 times larger. To maximize efficiency, we excluded loci where more than 45 SNPs fit this criterion with one exception, a block on chromosome 2 with 92 SNPs. This resulted in 1,198 SNPs in 64 schizophrenia loci moving forward to oligonucleotide design. Next we added SNPs from the AD GWAS (Lambert et al., 2013). Here we first used the lead SNPs reported by Lambert et al (Lambert et al., 2013) to identify all SNPs in strong LD (r^2^ > 0.9) using the bioinformatics tool SNAP (A. D. Johnson et al., 2008). The largest LD groups were then removed resulting in the addition of 30 SNPs across 9 AD GWAS loci to our assay, for a total of 1,228 SNPs selected to be included in the MPRA. All successfully assayed SNPs and the results of our assay are in supplementary Table 1.

### MPRA design and methods

The MPRA was designed to screen as many SNPs as possible for the cost of 1 oligo pool, with the understanding that the results are subject to future extensive validations by us or others. We performed 5 technical replicates, including each allele in 5 oligos with different barcodes (10 barcodes per SNP). To maximize the number of assayed SNPs, each allele was only profiled in one direction, since concordance between the two directions has been shown to be high (Klein et al., 2019).

Oligonucleotides of 150 bp including 95 bp centered on each SNP and 45 flanking based for amplification and cloning purposes (see below) were synthesized for each sequence of interest, as described in (Kheradpour et al., 2013) and (Melnikov et al., 2012). Target sequences were flanked with a multiple cloning site, a unique barcode and primers for PCR amplification (Figure 1). Each tested sequence (each allele in the case of SNPs) was tagged by 5 different barcodes, a number chosen to maximize the number of variants tested while still allowing the detection of significant outliers. We excluded any SNPs residing in genomic sequences such that the final oligo design would include restriction sites used for cloning (below). We successfully designed oligonucleotides for 1083 of our selected SNPs (1053 for SZ and 30 for AD), including 27 that had 3 alleles and 5 had 4 alleles.

**Figure 1:**
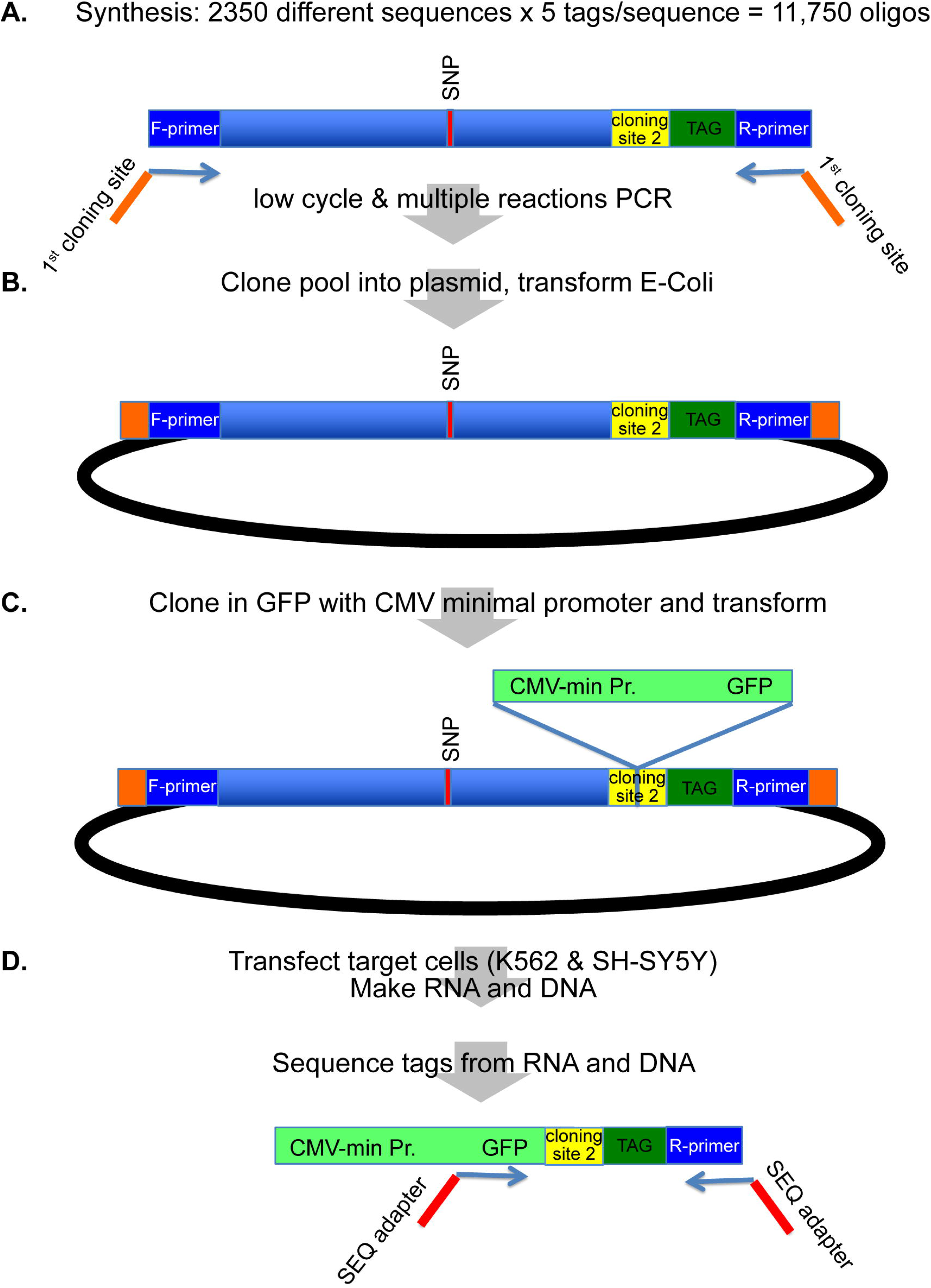
Experimental protocol for MPRA (See text for details).

We also included 545 oligonucleotides tiling the promoter of *EEF1A1* (chr6:73,520,057-73,521,235, hg38) in 95 bp segments with 10 bp overlap as a positive control. This is a gene with strong and widespread expression and our expectation was that some, but not all, of these sequences would strongly drive expression. A negative control was tiled segments of a 465 bp sequence from a pseudogene intron, overlapping no open chromatin signals in the encode data (chr1:14,992-15,456, hg38), which makes it less likely (but not impossible) to drive expression. The final pool of 11,750 uniquely barcoded oligonucleotides (see supplementary table 4) was ordered from the Broad Technology Labs (Broad Institute, Cambridge MA)

The library we received from the BROAD institute was amplified, cloned and transfected as described (Kheradpour et al., 2013; Melnikov et al., 2012) with slight modifications. The experimental design is summarized in Figure 1. First, using the primer sequences built into the synthetic oligos the library was amplified through a low-cycle (25 cycles) PCR reaction in 7 separate wells to preserve oligo representation and minimize PCR bias (Figure 1A). The 7 PCR products were gel-purified and pooled for subsequent steps. The primers used for PCR introduced two distinct SfiI sites as described (Melnikov et al., 2012). These sites were used for the subsequent directional cloning (1st cloning site in Figure 1A) into the pMPRA1 plasmid (Addgene ID# 49349, Figure 1B). The plasmids were used to transform E.Coli by electroporation. The synthesized oligos contained a second directional cloning site (restriction enzymes KpnII, XbaI) which was used in the next step to insert a CMV minimal promoter driving GFP between the putative regulatory sequence and the corresponding barcode sequence within the synthesized oligonucleotide (2nd cloning site, Figure 1C). After these two rounds of cloning - E.Coli transformation, we used the resulting plasmid library to perform three independent transfections of K562 (ATCC#CCL-243) chronic myelogenous leukemia cells and six independent transfections in SH-SY5Y (ATCC#CRL-2266) human neuroblastoma cells in two batches by lipofectamine3000 (Thermofisher Scientific cat. no L3000008). We then extracted DNA and RNA and performed 50 bp non-paired end sequencing reading from the 3’end, through the unique barcode and into the GFP transcript on an ILLUMINA MiSeq DNA Sequencer (Illumina, Inc. San Diego, CA). For each triplicate transfection we acquired 10 - 27 million reads of DNA and RNA.

All cell lines used for our experiments were donated from other Johns Hopkins Investigators and originally acquired from ATCC (www.atcc.org). The experimental process is summarized in Figure 1.

### Differential analysis

Before statistical analysis, we perform total count normalization for both RNA and DNA, that is we scale counts in each sample so that they all have the same library size. For differential analysis of MPRA activity levels between alleles, we use the mpralm method (Myint, Avramopoulos, Goff, & Hansen, 2019). This approach uses linear models to directly model activity measures and a combination of observation-level weighting and empirical Bayes methods to improve estimation of element-specific variances. The comparisons of activity measures between alleles are paired in the sense that each biological replicate measures the activity of all (2 or more) alleles of a SNP. The assaying of all alleles simultaneously in a given sample leads to correlation between their measurement, and for this reason, we use the mixed model approach of mpralm for making comparisons.

Our data for the SH-SY5Y cell line come from two separate batches, so before differential analysis we use the ComBat method (W. E. Johnson, Li, & Rabinovic, 2007) to correct the log-ratio activity measures. This method estimates location and scale parameters of batch effects at the element level and moderates these estimates using an empirical Bayes technique that pools information on these estimates across all elements. With these parameter estimates, batch-corrected outcome measures can be computed. From here, we proceed as detailed above with differential analysis.

### Barcode outlier analyses

To identify oligos that show outlier behavior in MPRA activity, we computed the coefficient of variation (CV) of RNA/DNA ratios across the 5 barcodes for a particular oligo. We report in Supplementary Table 1 two cutoffs on DNA counts to remove barcodes from the CV calculation if they were not sufficiently represented. We excluded barcodes with less than 5 and less than 10 DNA read counts. Of the SNPs that had at least one non-missing CV, we flag the SNPs that had at least one sample with a CV greater than the 99^th^ percentile threshold.

### Transcription factor binding analysis

To search for transcription factor (TF) motifs that can be affected by the SNPs in our study, we used the TFBSTools package available on Bioconductor (Tan & Lenhard, 2016). We used the union of the JASPAR2016 and ENCODE TF databases for a total of 2450 motifs. We only considered matches in which the motif overlapped the SNP position within the oligo and in which the position weight matrix (PWM) score was at least 70% of the maximum score possible for the PWM (i.e. a relative score of at least 70%) for at least one allele of a SNP.

### Comparisons with open chromatin measures

To assess whether MPRA active elements where overlapping with cell line-specific digital genomic footprinting (DGF) tracks we downloaded from UCSC the following files: hgdownload.cse.ucsc.edu/goldenpath/hg19/encodeDCC/wgEncodeUwDgf, file names wgEncodeUwDgfK562Hotspots.broadPeak.gz and wgEncodeUwDgfSknshraHotspots.broadPeak.gz. These correspond to data from K562 cells and retinoic acid treated SK-N-SH cells. SH-SY5Y are a sub-clone of SK-N-SH which is the closest cell line for which we could access DGF data.

## Results

### MPRA design and quality controls

#### Design

Details on the design are reported in the methods section. Briefly we selected from the reported SZ (Schizophrenia Working Group of the Psychiatric Genomics Consortium, 2014) and AD (Lambert et al., 2013) GWAS SNPs in high LD with the lead SNP at each locus, for a total of 1,053 SNPs in 64 SZ loci and 30 SNPs in 9 AD loci driven by the goal to screen as many loci as possible in one oligo pool. For each SNP we include 5 oligos (technical replicates) for each allele. To increase the number of assayed SNPs, each allele was only profiled in one direction as it has been shown that there is high concordance between the two directions (Klein et al., 2019). We synthesized a 95 bp region surrounding each allele.

As positive control we included sequences tiling the promoter of *EEF1A1* expecting that at least some (but not all) of these sequences will drive expression (see methods for details). As negative control we included sequences tiling a 465bp sequence from the intron of a pseudogene, not overlapping any open chromatin signal in Encode, suggesting – but not proving -- that this region does not drive expression.

#### Barcode representation

Of the initial 11,750 designed barcodes, 1,829 were never present in DNA or RNA, likely because of being absent or grossly underrepresented after initial library synthesis and later amplification (see methods). This left 10,106 barcodes (corresponding to 2,383 elements of the original 2,387) to be the total pool of possible barcodes for assessing experimental quality. Barcode representation (fraction of barcodes with non-zero counts) was high in both DNA and RNA in both tested cell lines (Table 1). For DNA, approximately 94% of barcodes were represented in K562 samples (where a sample is an independent transfection experiment) and SH-SY5Y batch 1 samples, and about 90% in SH-SY5Y batch 2 samples. For RNA, approximately 85% of barcodes were represented in K562 samples, 80% in SH-SY5Y batch 1 samples, and 60% in SH-SY5Y batch 2 samples. The lower representation in RNA is likely due to low/undetectable expression of some transfected oligonucleotides. All transfections included some unrepresented barcodes (Table 1, rows 3 and 4). After summing counts across barcodes for each element, only a small number of elements still had missing counts (Table 1, rows 5 and 6). We repeated these calculations when considering a barcode/element to be low or missing when the count was less than or equal to 2, and the resulting metrics are shown in the analogous rows in the bottom half of Table 1. We generally come to the same conclusions about the quality of barcode and element representation, but we do note that there is a noticeable increase in the fraction of barcodes that are always at a low count across all samples in SH-SY5Y batch 2. Since we pool count information across barcodes, this should not compromise our results, and we see that the number of MPRA elements that have low counts across all samples is still quite low. Further, although the majority of elements are represented by less than 5 barcodes, the majority of barcodes have average counts across samples that are sufficiently high (Supplemental Table 2.)

**Table 1.** Barcode and element representation across cell lines. This table is split into two sub-tables each with 6 rows. The top half of the table corresponds to a count threshold of 0, and the bottom half corresponds to a count threshold of 2. For both halves, the sample size for rows 1 to 4 is 10,106 barcodes, and the sample size for rows 5 and 6 is 2,383 elements. The ranges in rows 1 and 2 indicate a range over the 3 or 6 samples represented in a column.

|  | Threshold | K562 | SH-SY5Y (batch 1) | SH-SY5Y (batch 2) | SH-SY5Y (both batches) |
| --- | --- | --- | --- | --- | --- |
| Barcode represented > threshold: DNA | 0 | 94.7-95.6% | 93.7-96.9% | 87.3-92.1% | 87.3-96.9% |
| Barcode represented > threshold RNA | 0 | 84.9-85.5% | 78.5-81.0% | 57.1-81.7% | 57.1-81.7% |
| Barcodes always ≤ threshold DNA | 0 | 2.50% | 1.90% | 5.10% | 1.30% |
| Barcodes always ≤ threshold RNA | 0 | 7.90% | 13.10% | 15.30% | 9.10% |
| Elements always ≤ threshold DNA | 0 | 0.2% (5) | 0.3% (6) | 0.5% (11) | 0.2% (5) |
| Elements always ≤ threshold RNA | 0 | 0.4% (10) | 1% (24) | 1.2% (29) | 0.5% (12) |
| Barcode represented > threshold: DNA | 2 | 92.3-92.9% | 90.9-93.9% | 79.7-84.4% | 79.7-93.9% |
| Barcode represented > threshold RNA | 2 | 82.2-83.0% | 74.5-77.7% | 53.1-76.9% | 53.1-77.7% |
| Barcodes always ≤ threshold DNA | 2 | 4.80% | 4.30% | 11.70% | 4.10% |
| Barcodes always ≤ threshold RNA | 2 | 10.10% | 16.10% | 19.60% | 12.30% |
| Elements always ≤ threshold DNA | 2 | 0.4% (10) | 0.5% (12) | 0.9% (21) | 0.5% (11) |
| Elements always ≤ threshold RNA | 2 | 0.8% (19) | 1.4% (34) | 1.7% (41) | 0.9% (21) |

#### Correlation of count and activity measures between samples

To summarize count information for each MPRA element, we sum counts over barcodes and obtain one count per element per sample. These aggregated counts are used to compute activity measures

As we have previously shown this estimator is expected to have lower bias than an estimator that uses the mean of barcode-specific activity measures(Myint et al., 2019). Aggregated DNA counts show high between-sample correlations (Figures 2 and 3). Pairwise correlations of the log2-transformed aggregated counts ranged from 0.98 to 1.00. Correlations between samples within a single cell line were about the same as correlations between samples in different cell lines. For RNA, pairwise correlations of the log2-transformed aggregated counts ranged from 0.86 to 0.98 (Figure 2). Correlations between samples within a single cell line were somewhat higher than correlations between samples in different cell lines. The latter correlations ranged from 0.83 to 0.93.

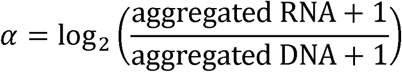

**Figure 2.**
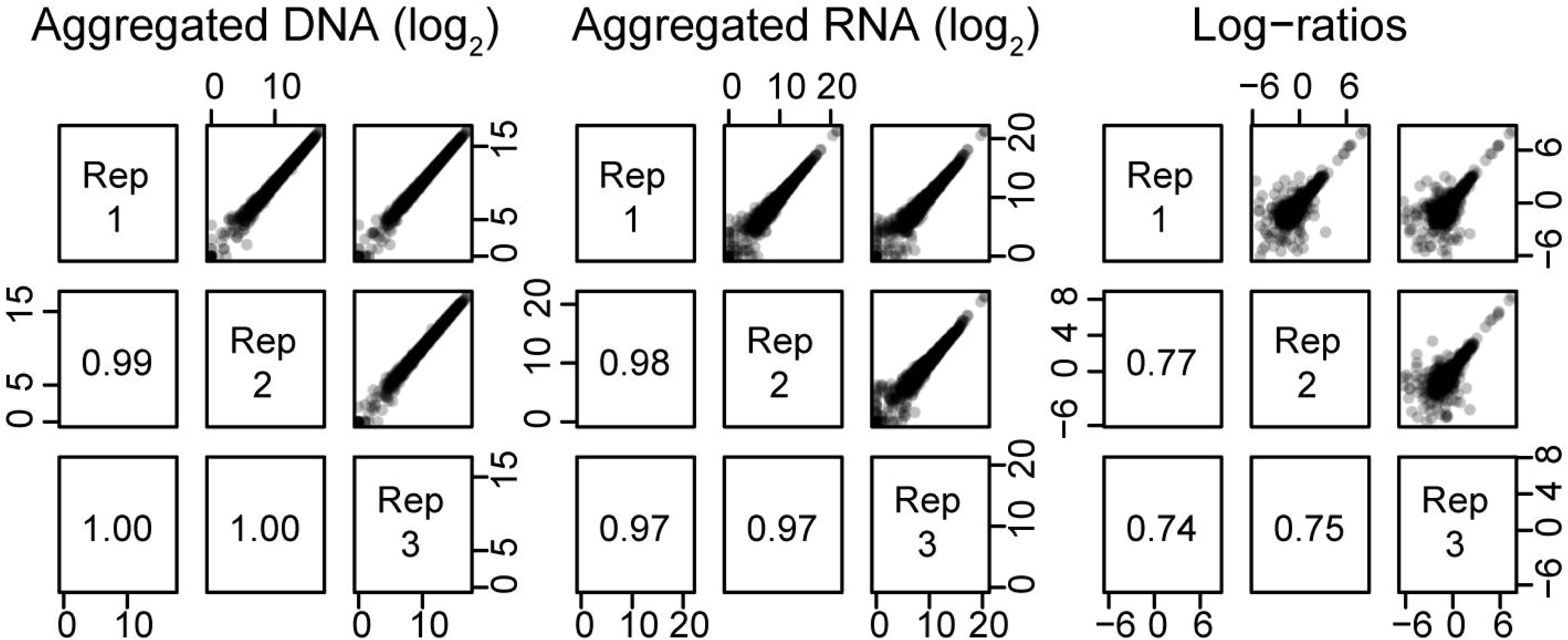
Correlations in the K562 cell line. The upper panels show scatterplots of MPRA quantities between samples. The lower panels indicate the correlation between a pair of samples.

**Figure 3.**
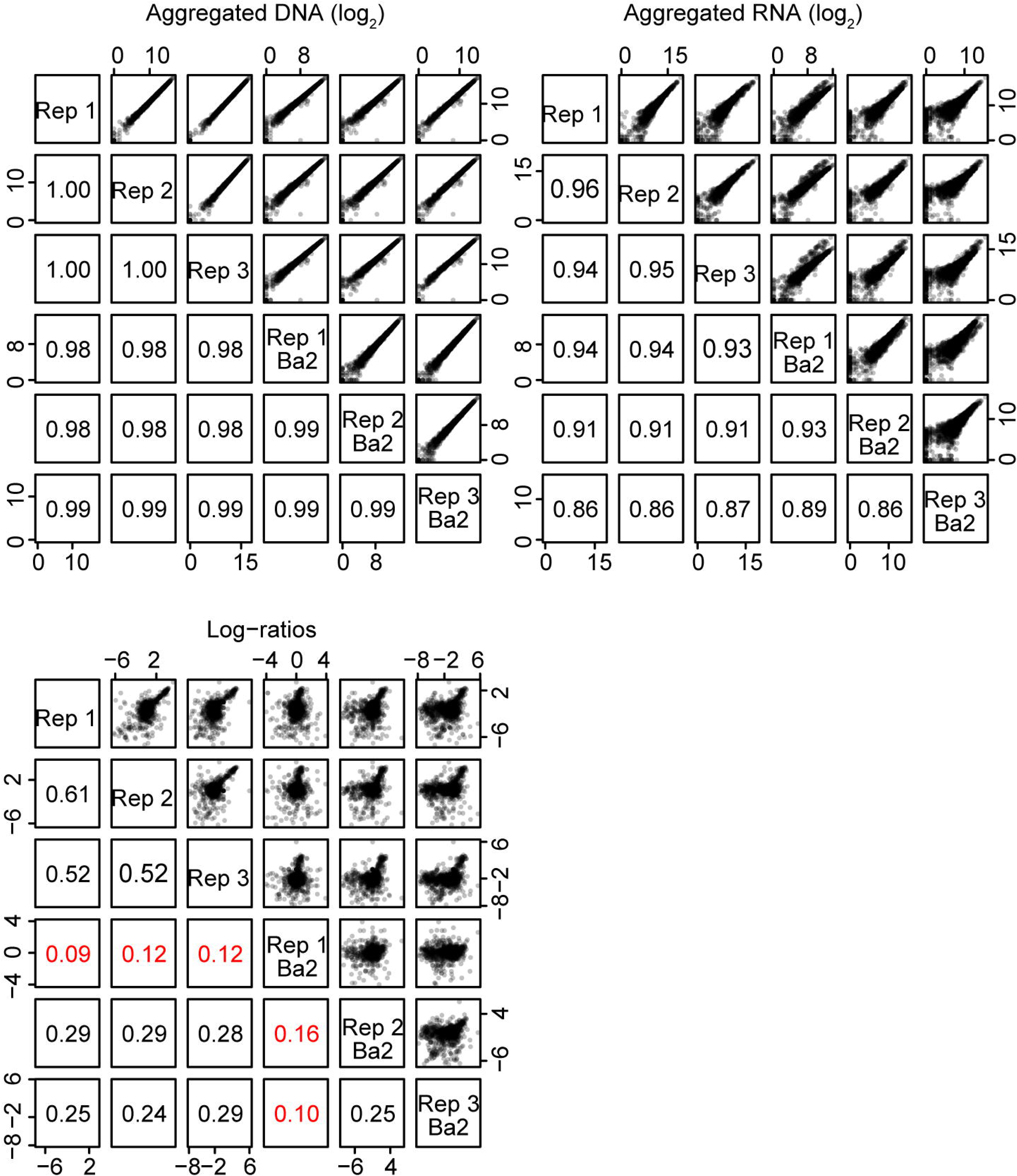
Correlations in the SH-SY5Y cell line. The upper panels show scatterplots of MPRA quantities between samples. The lower panels indicate the correlation between a pair of samples. Correlations are displayed in red if less than 0.2.

Activity measures (defined as α above) show between-sample correlations ranging from 0.74 to 0.77 in the K562 cell line (Figure 2). In the first batch of the SH-SY5Y cell line, activity measures show between-sample correlations ranging from 0.52 to 0.61 (Figure 3). In the second batch, between-sample correlations were low, ranging from 0.10 to 0.25. These correlation estimates include observations from elements without activity, and the second batch of SH-SYS5 include many such elements. When we remove activity measures with α < −1 (representing about 36% of activity measures α in the SH-SY5Y cell line), between-sample correlations for the second batch increase, ranging from 0.19 to 0.52. Generally, correlations between samples within a single cell line are higher than correlations between samples in different cell lines (Figure 4).

**Figure 4.**
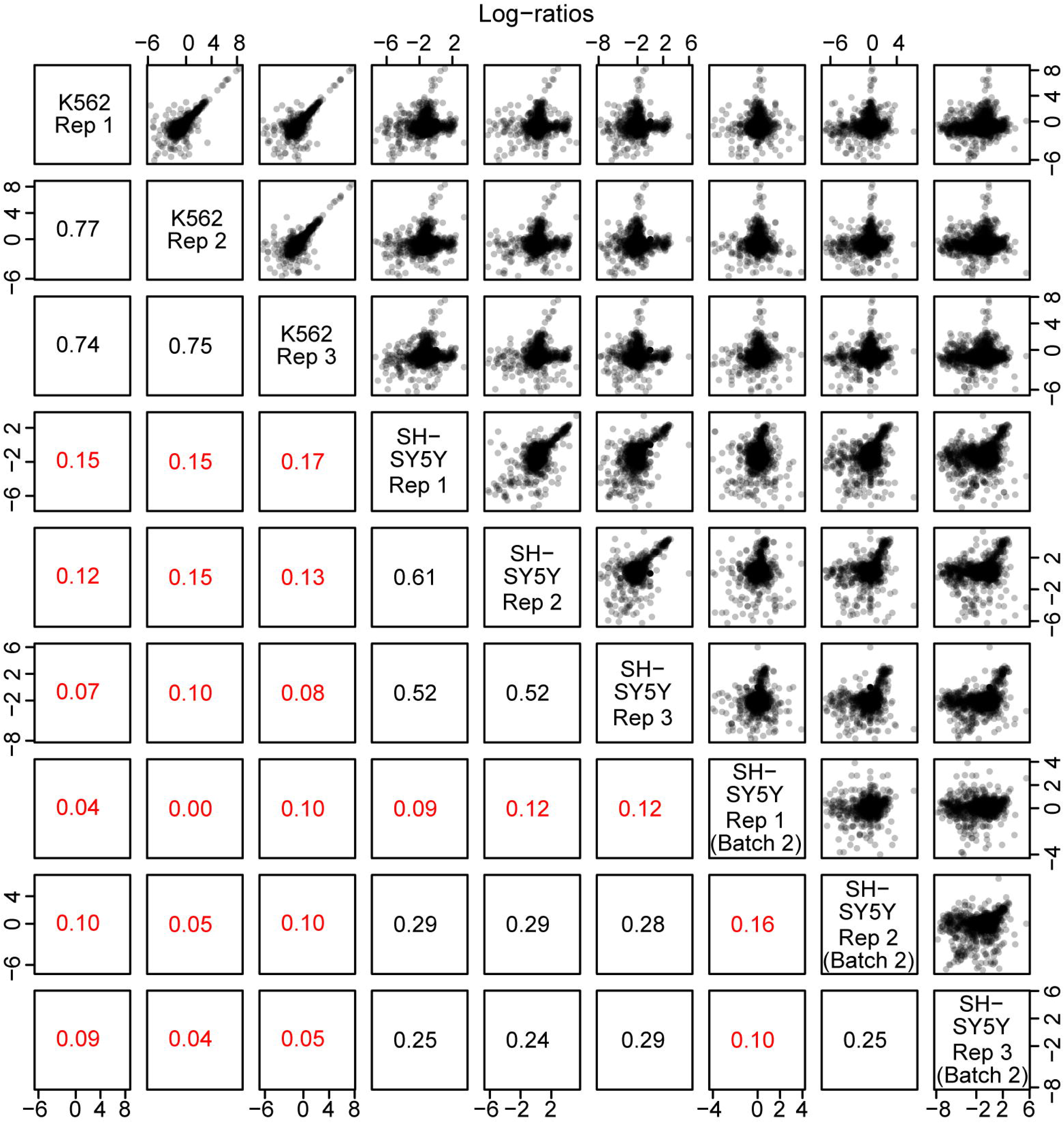
Correlations of log-ratio activity measures in both the K562 and SH-SY5Y cell lines. Correlations are displayed in red if less than 0.2. Generally, correlations within cell lines are higher than across cell lines.

We considered removing the second batch of SH-SY5Y cells from analysis because of the lower correlations observed for this experiment. In our experience, SH-SY5Y cells are more difficult to transfect and show lower transfection efficiencies compared to K562 cells, and we believe this is the cause of the lower correlations. However, despite the lower correlations, in our analysis of differential allelic activity (below) we found that including these samples lead to an increase in power, and we therefore decided to include them. Nevertheless, we present our analyses (below) with and without inclusion of this second batch (Supplementary Table 1)

#### Activity of positive and negative control sequences

We examined the positive control tiles from the positive control (*EEF1A1* promoter sequences) in the K562 cell line and observed that, as expected, many tiles showed sharp induction of reporter gene expression, where activity is defined as α above. Overall the *EEF1A1* promoter sequences (positive control) showed significantly higher activity than the negative control sequences (95% CI for difference in log-ratio activity measures: 0.50-1.12). While in the SH-SY5Y cell line all positive control sequences together did not show significantly higher activity, the same sequences that were active in K562 were also significantly active. Figure 5 demonstrates this across all thresholds for defining high activity. When considered as strong positives the tiles with activity levels α above 2 in K562, the same tiles consistently show higher expression than the negative controls in SH-SY5Y. Not surprisingly given its overall better performance, batch 1 shows significant increases in positive control activity over a wider range of thresholds than batch 2 (Figure 5).

**Figure 5.**
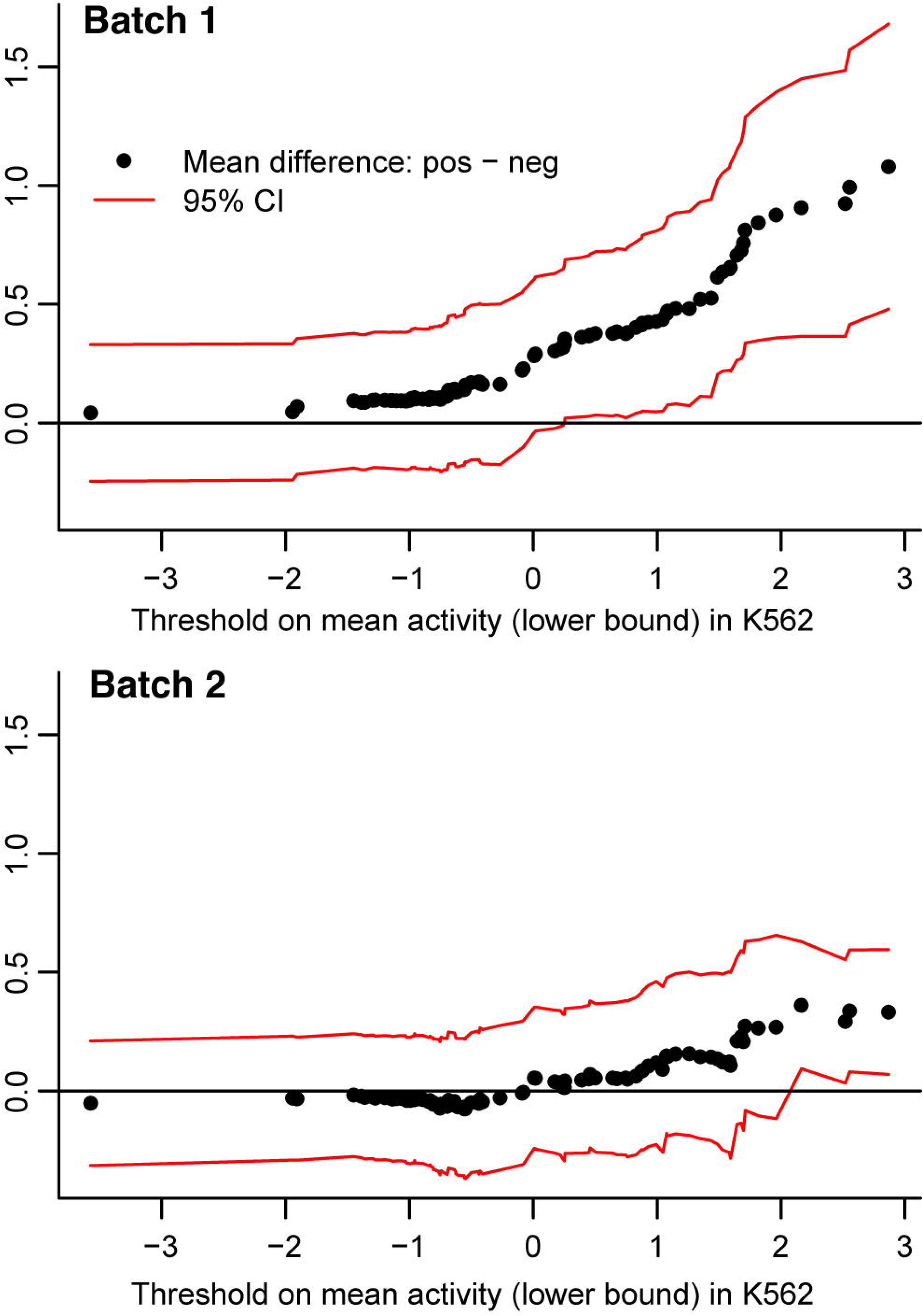
Comparison of activity levels in subsets of *EEF1A1* positive control and negative control sequences: neuronal cell line. The x-axis shows different activity thresholds for defining the set of positive controls, using data in the K562 cell line. The set of negative controls is fixed. For each set of positive controls, we show the difference with the negative controls (Y axis) and the confidence intervals for the difference. In batch 1, moderately active positive control oligos (K562 mean activity greater than ∼ 0) show significantly higher activity than the negative controls. In batch 2, only the most highly active positive control oligos show significantly higher activity than negative controls.

### Evaluation of variants that show differential enhancer activity

#### Barcode outlier analyses

To assess whether some tested elements might be less reliable due to specific outlier barcode counts, and because standard approaches such as calculating the number of standard deviations away from the median cannot be applied to samples with 3 or 4 barcodes present, we calculated the coefficient of variation of counts for each MPRA element. This measure of dispersion provides a metric where higher values signify less confidence. To avoid inflated ratios due to small denominators we only included barcodes with at least 5 DNA reads. Barcodes with fewer reads would not significantly influence the final calculations as these were made with aggregated counts and small numbers would be of little consequence. Setting a cutoff at the 99th percentile of the distribution we found 0.4% (4/1072) of the barcode sets for K562, 2.0% (21/1068) for SH-SY5Y-batch-1, 3.2% (33/1047) for SH-SY5Y-batch-2 and 5.1% (54/1068) for SH-SY5Y-both-batches to contain outliers. All individual CVs for each allele are listed in Supplemental Table 1 so each SNP can be individually evaluated.

#### Location of SNPs with differential enhancer activity

We successfully assayed 1,079 SNPs and found 144 SZ and 4 AD SNPs that show differential enhancer activity between alleles (significant allelic skew) in the K562 cell line and 50 and 3 in the SH-SY5Y cell line (FDR < 0.05). These SNPs, which we here call significant SNPs, are located on several different chromosomes (Figure 6). Nine SNPs showed allelic differences in both K562 and SH-SY5Y: rs73036086, rs2439202, rs6801235, rs134873, rs13250438, rs2605039, rs7582536, rs1658810, rs8061552. Of those all except rs2439202 were in the same direction (binomial p=0.02). Six of the remaining 8 (Table 2) were also eQTLs for one or more gene in the dorsolateral prefrontal cortex (DLPFC) in the CommonMind consortium data (http://www.nimhgenetics.org/available_data/commonmind). Four of those were also eQTLs for one or more of the same genes in at least one tissue in the GTEx database (https://gtexportal.org) while rs6801235 was an eQTL for a different gene in GTEx, in the cerebellar hemisphere (see last two columns of Table 2). Note that the sample sizes of brain related tissue collections in GTEx are generally around 150, four times fewer than the CommonMind DLPFC collection.

**Figure 6.**
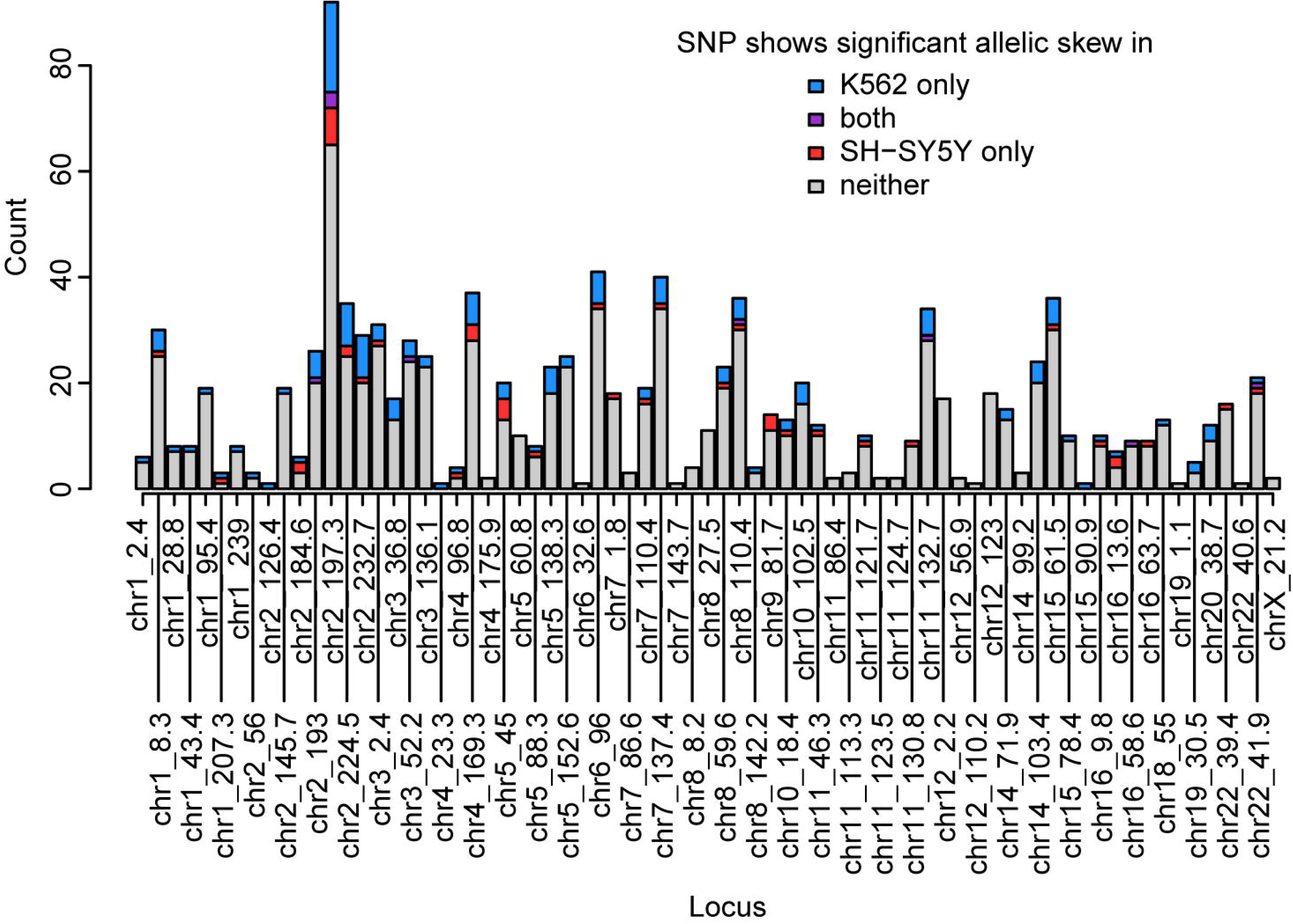
Genomic locations of assayed SNPs and SNPs showing allele specific activity. The SNPs assayed in this MPRA come from a variety of linkage disequilibrium (LD) blocks across several chromosomes. SNPs that show differential regulatory activity between alleles are also present in a wide variety of loci. Loci are labeled by chromosome and location on hg19.

**Table 2.**
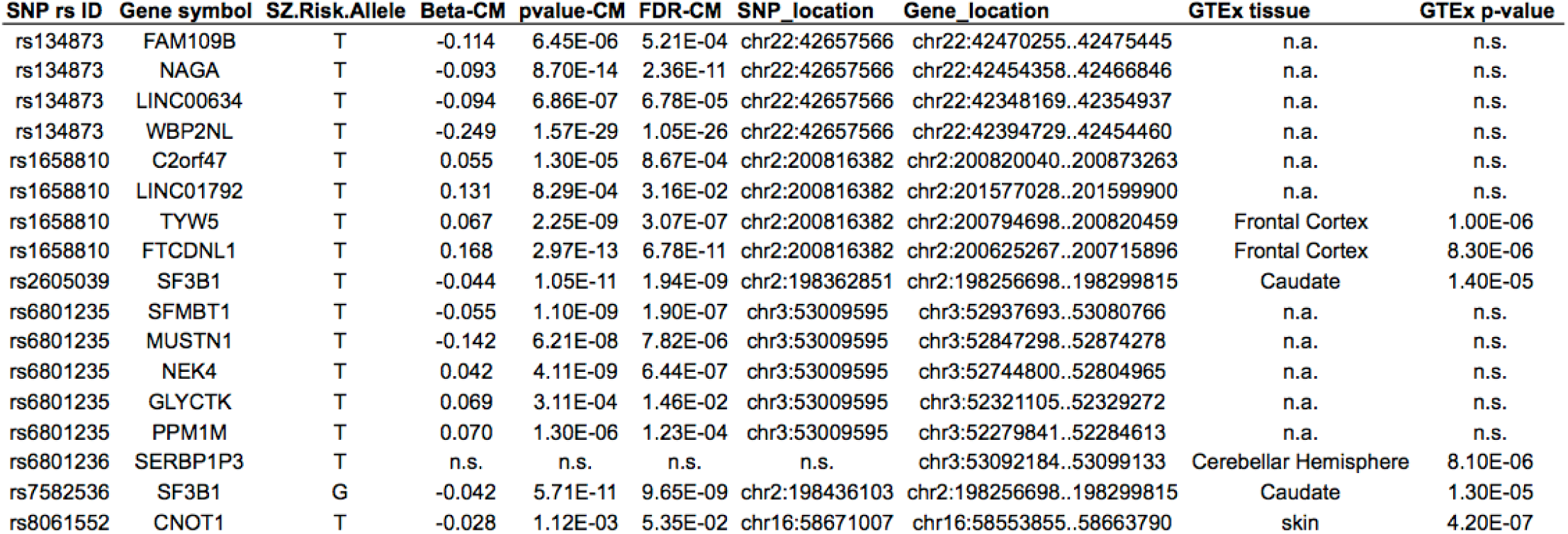
EQTLs among the 8 SNPs significant in both cell lines. The rs ID from dbSNP and the gene symbol are followed by the SNP risk allele (from the PGC GWAS)

On average, there are 2.6 significant SNPs per locus (median: 1 significant SNP per locus), and on average, 18% of SNPs tested were significant (median: 13%). Out of the 73 successfully tested loci, 19 do not have any significant SNPs, and 5 of these 19 are single-SNP loci. Of the 54 GWAS loci with SNPs showing significant allelic differences, in 31 there were eQTLs for nearby genes among the SNPs (66%), however the rate was also high for 16 of the loci not showing significant allelic differences (74%).

#### There is no preferred direction of effect on enhancer/repressor activity for risk versus non-risk alleles

If a SNP changes SZ or AD risk by affecting a nearby gene enhancer, is the risk allele more likely to decrease or increase enhancer activity? In other words, are risk alleles more often associated with loss or gain of enhancer function? To explore this question, we overlaid our MPRA results with GWAS information on risk effect directions (i.e. which allele increases expression vs. which allele increases risk). Because of the larger number of SNPs and recognizing that this might be different between diseases we did the analysis only for SZ SNPs. Of the 144 significant SZ SNPs in K562, 81 are SNPs for which the risk allele from GWAS shows lower MPRA enhancer activity than the non-risk allele (56%, 95% binomial CI: 47%-64%). Of the 50 significant SZ SNPs in SH-SY5Y 25 are SNPs for which the risk allele shows lower MPRA enhancer activity (50%, 95% binomial CI: 35%-65%). Overall, these results suggest no specific direction of effect on expression characterizing disease risk variants. It must be noted however that this analysis examines the effect of each individual SNP on the disease-associated haplotype in isolation. It is likely that selection has favored alleles with opposing effects to be on the same haplotype, balancing each other towards a more favorable total combined effect where direction might be more consistent across genes. Such interactions would not be captured in our experiment.

#### MPRA activity levels show concordance with chromatin accessibility measures

We find that MPRA activity levels in the two cell lines show good overlap with cell line-specific digital genomic footprinting (DGF) tracks from UCSC (Methods). In particular, highly active SNPs in the K562 cell line show greater enrichment for K562 DGF sites than highly active SNPs in the SH-SY-5Y cell line. Inversely, highly active SNPs in the SH-SY-5Y cell line show greater enrichment for SkNSHRA (a retinoic acid-treated cell line of which SH-SY5Y are a sub clone) DGF sites than highly active SNPs in the K562 cell line (Figure 7). We also considered overlaps with specific enhancer chromatin markers, however these were too sparse for meaningful comparisons. For example the K562 H3k27ac sites overlap only about 4.9% of non-significant SNPs and 5.4% of K562 significant SNPs, which is only 8 SNPs.

**Figure 7.**
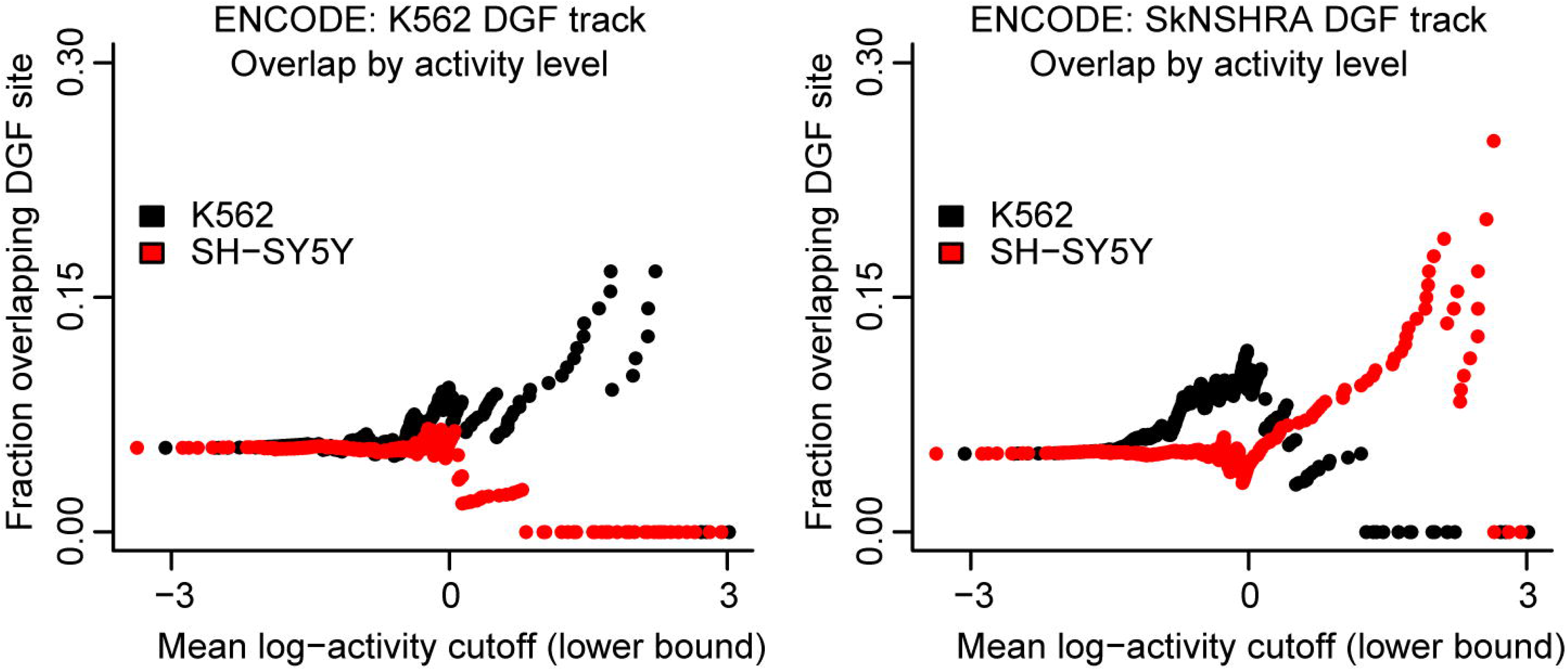
DGF enrichment by activity level. Every point corresponds to a set of SNPs. This set is defined by a cutoff on the mean MPRA activity. For these sets, the fraction of SNPs overlapping a DGF site is shown on the y-axis for the K562 cell line (left) and for the SkNSHRA cell line (right).

#### Identification of disrupted features contributing to differential activity

We searched for transcription factor (TF) binding motifs overlapping SNP positions and compared binding scores between alleles to assess potential disruptions in TF binding. We used the union of the JASPAR2016 and ENCODE databases for a total of 2450 TF motifs. The main match metric we use fro every position is the relative score, the ratio of the observed position weight matrix (PWM) score divided by the maximum possible PWM score, giving relative scores between 0 and 1. Due to the large number of motifs examined, most SNPs in our MPRA assay showed high relative score for at least one TF motif. In fact, all MPRA SNPs have at least one TF motif match with a relative score of at least 0.963 (Figure 8).

**Figure 8.**
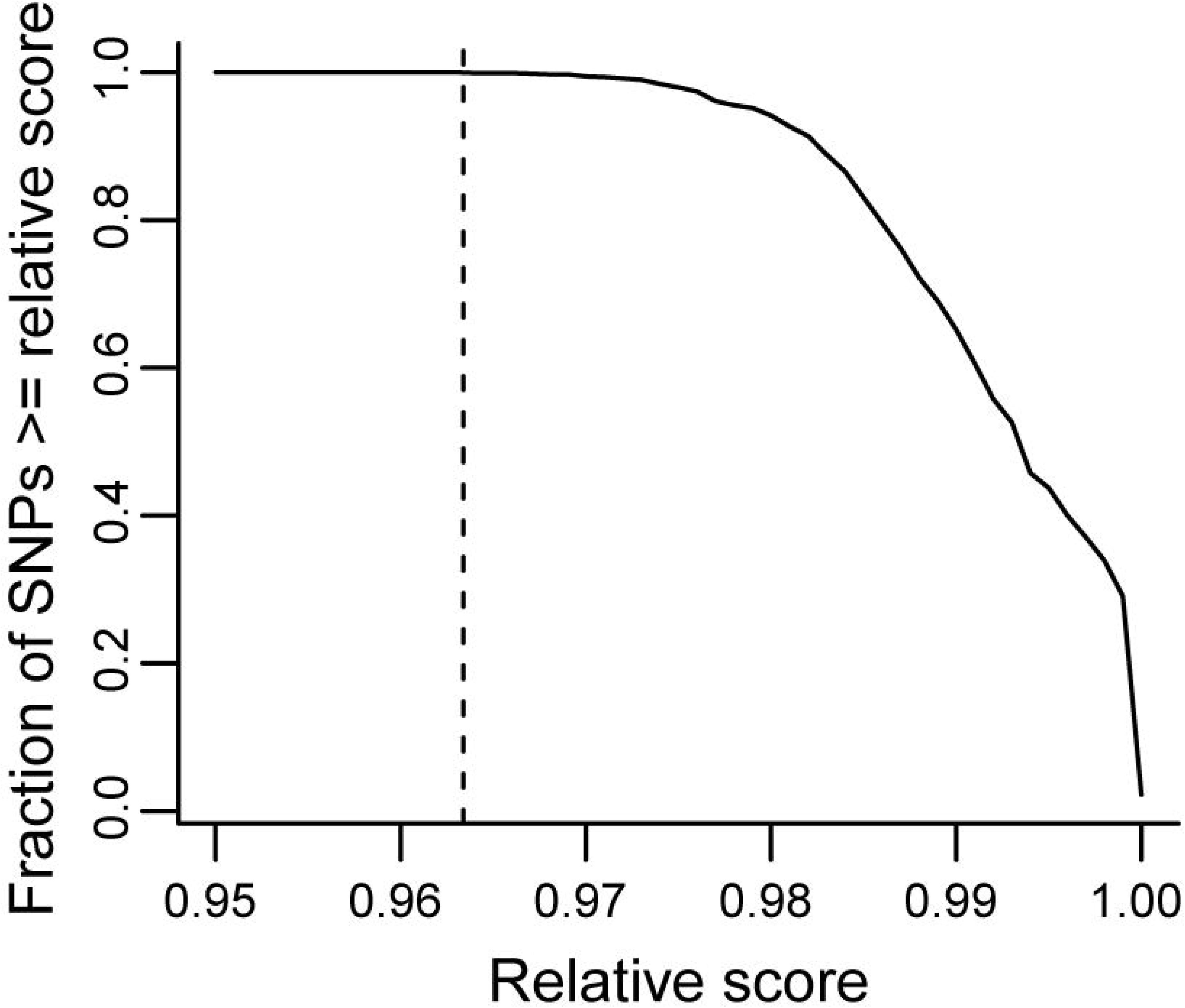
Distribution of PWM relative scores. The y-axis shows the fraction of MPRA SNPs that have at least one TF motif match with a relative score greater than or equal to the value given on the x-axis. The dashed vertical line shows the highest relative score (0.963) at which all SNPs still have a motif match.

Given that a PWM relative score gives a measure of how strongly a particular TF might bind to a region, we asked if a higher score was associated with higher enhancer activity in the MPRA. For each match between a TF motif and an MPRA oligo, we obtained a PWM relative score for each allele and the mean activity from the MPRA data and determined whether they were correlated. We did not observe such a correlation in either the K562 or the SH-SY5Y cell line. This is probably expected given the noise in examining such a wide variety of motifs across so many sequences. We do see that all MPRA SNPs with significant allelic differences in activity have at least one TF motif that shows disruption in binding in agreement with MPRA activity levels, but it is unclear whether this is meaningful since MPRA SNPs with non-significant allelic differences show the same.

#### Support for combinatorial SNP effects leading to large LD blocks

Ours (Eckart et al., 2016) and others’ work (Corradin et al., 2014) suggests that within large LD blocks there are often multiple regulatory variants that presumably combine their effects in regulating their target gene. Work on Mendelian disorders has also shown that favorable combinations of alleles can become more frequent due to selection (Castel et al., 2018). We hypothesize when there are more than one functional polymorphic sites in a region, selection might favor specific combinations of alleles, bringing the expression of a gene closer to optimal. Such selection will generate groups of SNPs in LD all of which will be associated with disease and many will be functional. The observable outcome would be that larger groups of SNPs associated with disease will have a higher density of functional SNPs than smaller groups, where this mechanism has not influenced LD. Our selection strategy where lead SNPs were selected based on p-value and the remaining SNPs based on the fold difference in p-value from the lead avoids any bias for stronger signals to include more SNPS, and the group size only depends on the LD structure. Also, since we have a mix of smaller and larger LD groups this provides an opportunity to test this hypothesis. We did so on our K562 cell line data, as it had the most positive results and to avoid confounding from differences in regulation between cell lines. Our selection strategy makes smaller blocks when there are fewer SNPs in LD with the lead SNP, regardless of the lead SNP p-value. Since the lead SNP also has the highest probability to be functional this biases smaller blocks towards higher densities of positives. To reduce this bias without introducing other biases we removed the lead SNP from all blocks before this analysis. We also remove the largest LD block of 92 SNPs as an outlier from other blocks (although we note that including it improves the significance of the results). We then tested whether there is a correlation across loci between block size and the fraction of positives 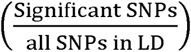 in each block. Figure 9 shows the scatterplot and best fit line showing a positive correlation significant at p=0.04 (two sided). Although the significance is modest it supports the hypothesis that combinatorial SNP effects lead to larger LD blocks.

**Figure 9.**
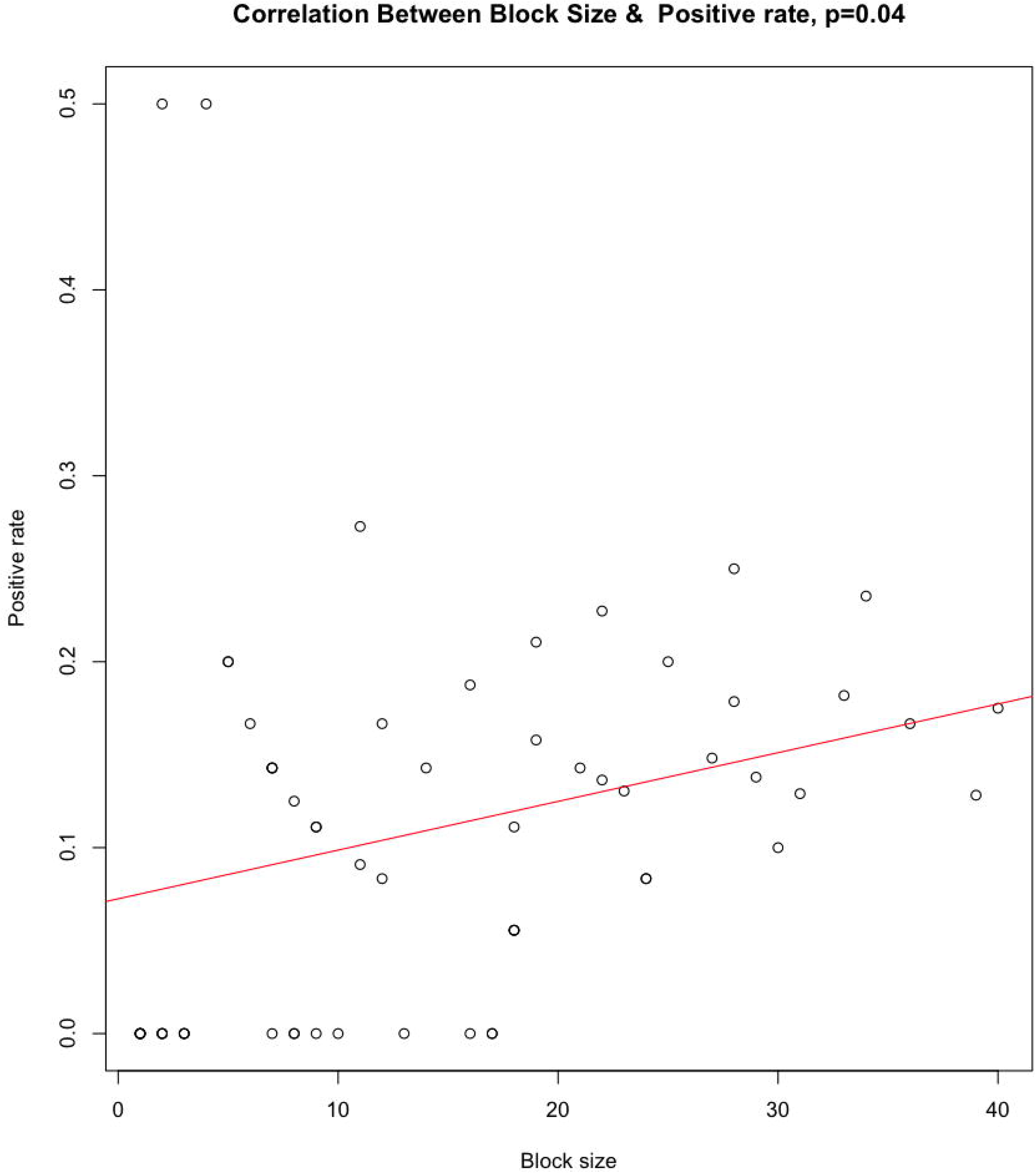
Scatterplot of block size by fraction of positives in the block. We observed a positive correlation (p=0.04) which supports prior observations that combinatorial SNP effects on gene expression may drive selection of haplotypes and larger LD blocks

## Discussion

We report the results of a screen for functional SNPs associated with SZ and AD on two different cell lines, K562 and SH-SY5Y. The purpose of this work is to add to the data available to us and other investigators in the effort to prioritize for functional follow up among the multiple SNPs showing statistical associations with disease. We report a total of 192 SNPs among the total of 1079 tested (18%) that show statistically significant allelic differences (FDR = 0.05) in driving reporter gene expression. While there are limitations inherent to such experiments as we discuss, these variant sequences are good candidates for more detailed functional follow up beyond reporter assays when investigating disease associations and will be an important resource for SZ and AD research.

We made a number of observations that support the validity of our results. The positive and negative controls behaved as expected and the concordance of direction of signals that emerged as significant in both cell types suggest that we are mostly looking at true signals. The significant and cell type specific overlap with open chromatin marks further supports the validity. Whether a higher overlap should be expected between the two cell lines is less clear, given significant regulatory differences between cell lines as suggested by the also limited overlap of DGF hotspots. Based on the encode datasets described in the methods we find that only 19.7% of the K562 DGF hotspots show any overlap with SK-N-SH hotspots.

We observe a high number of eQTLs, at ∼70% of tested loci, though it is not different between loci harboring detected significant SNPs and those that do not. It is not clear whether we should expect such a difference, given the differences between the cell lines we used and the DLPFC bulk tissue that was used to identify eQTLs and the very likely presence of false negatives in our results. Of the genes regulated by the eQTLs shown in Table 2 four are of particular interest as they are supported in both CommonMind and GTEx datasets. *TYW5* encodes a protein involved in tRNA modification and is essential for translational fidelity (Kato et al., 2011). It is among the genes reported to show significant transcript abundance change from prenatal to postnatal supporting a role in brain development (Birnbaum et al., 2015). *FTCDNL1* on the other hand which is at the same locus with *TYW5*, is a gene for which very little is known. *SF3B1* encodes a splicing regulator (Kfir et al., 2015) and has also been implicated in SZ through expression analysis in a rat psychosis model (Ingason et al., 2015). Interestingly changes in splicing have also been associated with SZ (Morikawa & Manabe, 2010; Wu et al., 2012). Like *TYW5, SF3B1* also shows transcript abundance changes from prenatal to postnatal time (Birnbaum et al., 2015). Finally, *CNOT1* encodes an important subunit for the enzymatic activity of the CCR4-NOT complex, critical for mRNA deadenylation and decay. This gene has been reported to participate in a network of 12 SZ-associated genes with correlated expression.(Liedtke, Zhang, Thompson, Sillau, & Gault, 2017).

Another interesting result is the correlation between the size of an LD block (in terms of number of SNPs) and the fraction of positives within it. While marginally significant this is consistent with many previous observations that suggest that there might be combinatorial effects between many functional variant sequences in each haplotype and that such haplotypes may be under selection. Such a phenomenon is very important to be aware of as we enter a time when many laboratories are focusing on understanding the biological consequences of variants revealed by large GWAS.

When considering our results, it is important to remember that the caveats affecting classic reporter assays, an artificial expression system, also apply here. The frequency of false negatives is likely high as this expression system only poorly mimics that of native genes. False positives are also possible, for example if the regulatory effect on the reporter gene comes from other parts of the construct or if in the disease-relevant tissues the tested variant resides in closed chromatin. Having tested two cell types reduces false negatives, but it might also somewhat increase these types of false positives. It is also important to note that the results from the SH-SY5Y cells were significantly weaker than those of the K562 cells. Our experience has been that SH-SY5Y cells are consistently less efficient for transfection both in these and in other experiment, which explains the lower oligo representation and lower yield of positive results. While we included SH-SY5Y because as a neuroblastoma cell line might be better for studying brain disorders, we should note that, in fact, K562 might also be quite relevant. For one, it is enriched for genes expressed in immunity-related tissues reported by the PGC for the SZ GWAS signals (Lambert et al., 2013). We have also observed that the overlap of chromatin marks and SZ SNPs is higher for K562 cells than SK-N-SH-RA that SH-SY5Y are derived from (5.4% of all SZ SNPs we assayed overlap K562 DGF hotspots compared to 5.1% overlapping SK-N-SH-RA DGF hotspots).

Our results provide an initial screen for functional elements. More labor-intensive methods would need to be employed for each one of these variants to confirm their function. Such methods, as for example specific genomic editing of variant bases and examination of the consequences in disease relevant brain cell types, can provide a definite answer regarding their role in disease.

## Acknowledgements

This work was supported in part by NIMH grants R56MH113215 to DA and P50MH094268 (project 1 PI: DA).

## Supporting Information Legends

**Supplementary Table 1:** This excel file includes two tabs. The tab “ALL_SNP_results” lists all 1,079 SNPs successfully assayed in this study along with their location, uncorrected and corrected p-values for allelic differences and the disease which they are associated with. The tab “Only with SH-SY5Y batch 2” Includes the names and locations of SNPs that only reach statistical significance (corrected) when both batched of transfections to SH-SY5Y were included (see text for details).

**Supplementary Table 2:** Breakdown of barcode representation. The sample size for the first 8 rows is 2,387 MPRA elements. The sample size for the last 2 rows is 11,935 barcodes.

**Supplementary Table 3:** Sequences of all synthesized oligonucleotides

