## Supplementary figures and images for "A screen of 1,049 schizophrenia and 30 Alzheimer’s-associated variants for regulatory potential"

### SUPPLEMENTAL FIG1

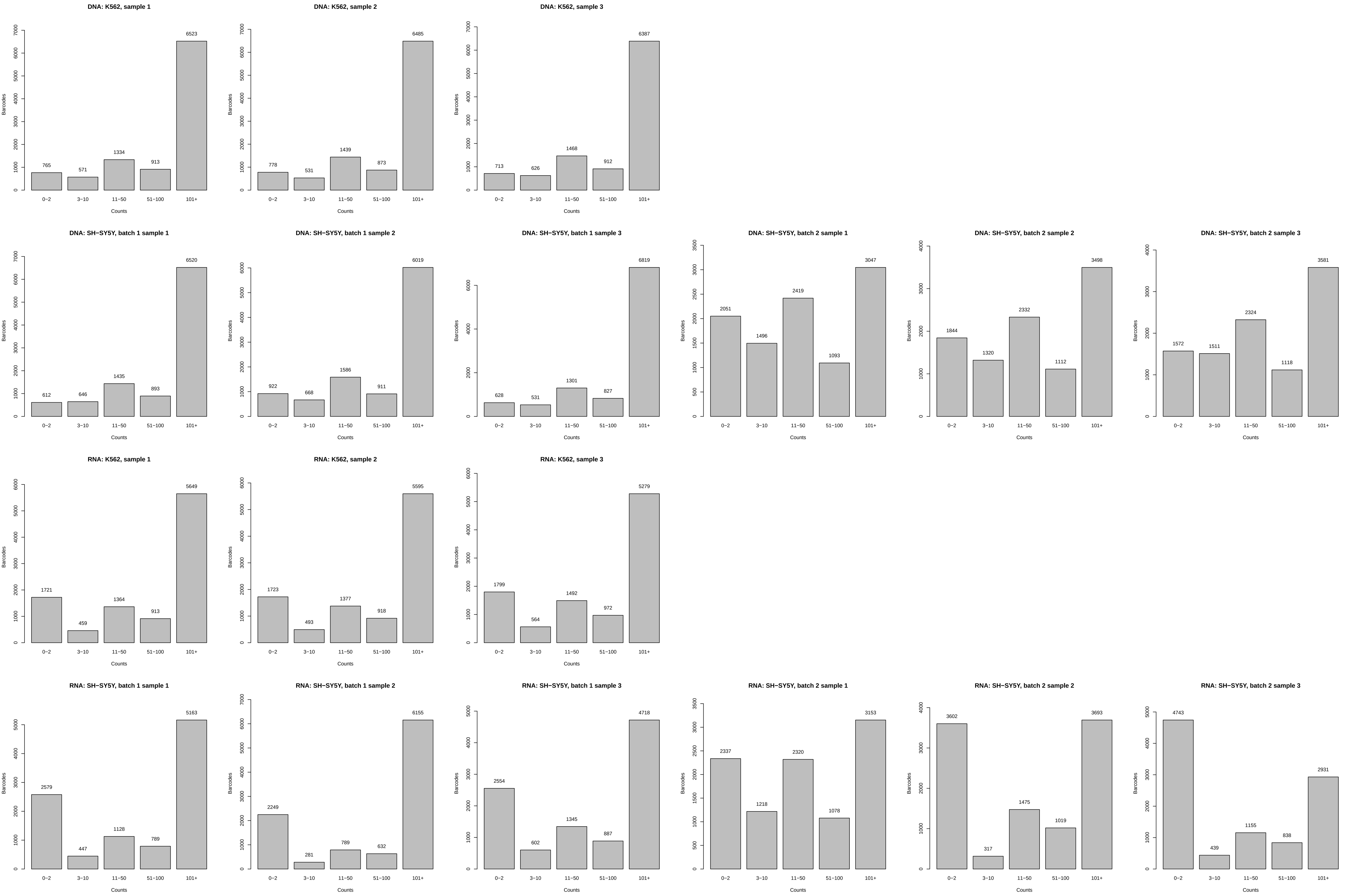
