## SUPPLEMENTAL TABLE 2 for "A screen of 1,049 schizophrenia and 30 Alzheimer’s-associated variants for regulatory potential"

|  | K562 | SH-SY5Y (batch 1) | SH-SY5Y (batch 2) | SH-SY5Y (both batches) |
| --- | --- | --- | --- | --- |
| Elements represented by less than 5 barcodes in all samples: DNA | 58.4% | 57.1% | 63.3% | 56.1% |
| Elements represented by less than 5 barcodes in all samples: RNA | 68.3% | 75.3% | 77.3% | 71.3% |
| Elements represented by less than 4 barcodes in all samples: DNA | 21.3% | 20.5% | 25.8% | 19.9% |
| Elements represented by less than 4 barcodes in all samples: RNA | 31.5% | 39.7% | 43.7% | 35.8% |
| Elements represented by less than 3 barcodes in all samples: DNA | 6.2% | 5.6% | 8.4% | 5.4% |
| Elements represented by less than 3 barcodes in all samples: RNA | 11.5% | 15.6% | 19.1% | 13.1% |
| Elements represented by less than 2 barcodes in all samples: DNA | 1.7% | 1.6% | 2.6% | 1.6% |
| Elements represented by less than 2 barcodes in all samples: RNA | 3.1% | 5.5% | 5.9% | 4.0% |
| Fraction of barcodes where geometric mean count over samples is at least 10: DNA | 94.7% | 94.5% | 58.3% | 86.4% |
| Fraction of barcodes where geometric mean count over samples is at least 5: RNA | 97.4% | 96.9% | 47.6% | 90.2% |

**Supplemental Table 2:** The sample size for the first 8 rows is 2,387 MPRA elements. The sample size for the last 2 rows is 11,935 barcodes.
